# Detection of M-Protein in Acetonitrile Precipitates of Serum using MALDI-TOF Mass Spectrometry

**DOI:** 10.1101/2020.03.13.990192

**Authors:** Nikita Mehra, Gopal Gopisetty, S Jayavelu, Rajamanickam Arivazhagan, Shirley Sundersingh, Parathan Karunakaran, Jayachandran Perumal Kalaiyarasi, Krishnarathinam Kannan, Venkatraman Radhakrishnan, Tenali Gnana Sagar, Thangarajan Rajkumar

## Abstract

**Purpose of the research:** Multiple myeloma and plasmacytomas belong to a group of disorders, namely plasma cell dyscrasias and are identified by the presence of a monoclonal protein (M-protein). MALDI-TOF-mass spectrometry (MS) has demonstrated superior analytical sensitivity for the detection of M-protein and is now used for screening of M-protein at some centres. We present the results of an alternative methodology for M-protein analysis by MALDI-TOF MS.

**Methods:** Serum samples from patients with newly diagnosed multiple myeloma or plasmacytoma with positive M-protein detected by serum protein electrophoresis, immunofixation electrophoresis and serum free light chain analysis, underwent direct reagent-based extraction process using Acetonitrile (ACN) precipitation. Serum *κ* and *λ* light chains were validated using immunoenrichment by anti- *κ* and anti- *λ* biotin-labelled antibodies immobilised on streptavidin magnetic beads. MALDI-TOF MS measurements were obtained for intact proteins using alpha-cyano-4-hydroxycinnamic acid as matrix. The images obtained were overlaid on apparently healthy donor serum samples to confirm the presence of M-protein.

**Principle results:** Characteristic M-protein peaks were observed in the ACN precipitates of serum in the predicted mass ranges for *κ* and *λ*. The *κ* and *λ* peaks were confirmed by immunoenrichment analysis. Twenty-seven patient samples with either newly diagnosed multiple myeloma or plasmacytoma with monoclonal gammopathy detected by the standard methods were chosen for Acetonitrile precipitation and analysed by MALDI-TOF MS. All 27 patient samples demonstrated a peak suggestive of M-protein with mass/charge *(m/z)* falling within the *κ* and *λ* range. The concordance rate with serum immunofixation electrophoresis and free light chain analysis was above 90%.

**Major conclusions:** We report the results of a low-cost reagent-based extraction process using Acetonitrile precipitation to enrich for *κ* and *λ* light chains, which can be used for the screening and qualitative analysis of M-protein.

## 1.1 Introduction

Multiple myeloma (MM) and plasmacytoma(s) belong to a group of clonal plasma cell dyscrasias. (1,2) The diagnosis of these disorders is established by the detection of abnormally secreted monoclonal immunoglobulin-M-protein in the blood and sometimes in the urine. Around 2-3% have truly non-secretory multiple myeloma, i.e., absence of M-protein. (3)

M-protein is analysed by a multitude of tests, including serum/ urine protein electrophoresis (SPEP/ UPEP), serum/ urine immunofixation (IFE) and serum free light chains (sFLC). The combination of these tests can be laborious, time-consuming and expensive. (4,5) Recent studies have demonstrated the feasibility of employing matrix-assisted laser desorption ionization mass spectrometry-time of flight mass spectrometry (MALDI-TOF MS) for the detection of M-protein in MM and AL amyloidosis termed MASS-FIX. (6–10) MASS-FIX has demonstrated superior analytical sensitivity to IFE and is now used in the clinic. (11) MASS-FIX can be utilised for rapid and accurate estimation of M-protein for screening and diagnosing plasma cell disorders.

The advantage of using MALDI analysis is based on the ability to measure the mass to charge *(m/z)* ratios of the light chains: *κ m/z* ([M+2H]^2+^: 11550-12300 Da, [M+H]^+^: 23100-24600 Da), and *λ m/z* ([M+2H]^2+^: 11100-11500 Da, [M+H]^+^: 22200-23100 Da) and high throughput automated analysis. (7) In a healthy individual, secretion of the immunoglobulins by plasma cells in the bone marrow is polyclonal with several plasma cells secreting these proteins. The polyclonal nature of the secretion, upon the analysis in a mass spectrometer, would give rise to a spectrum indicating a Gaussian distribution pattern with no single protein clone (*κ* or *λ*) being predominant. (12) However, in the diseased state, due to increased secretion of antibodies by a few clones, there is a distinct secretion pattern which is contrary to the Gaussian distribution pattern and is reflected in MALDI-TOF MS analysis. This distinction in the pattern of protein profile between the healthy and diseased state is diagnostic for monoclonal gammopathy. Another necessary consequence of the analysis of antibody light chains using MALDI-TOF, is that the *m/z* measurements obtained from the instrument help to determine the nature of the disease which can be categorized as either *κ* or *λ*. MASS-FIX is a mass spectrometry-based analysis of M-protein which entails isotype-specific nanobody enrichment along with MALDI-TOF MS. A significant drawback of this procedure is the high cost of the reagents involved. Additionally, the primary reagent required, namely the *κ* and *λ* specific antibodies, are qualitatively sensitive and prone to show variation during production.

Therefore, we felt the need to device a low-cost reagent-based extraction process which can enrich for the *κ* and *λ* proteins and is simultaneously compatible with MALDI-TOF MS.

## 1.2 Materials and Methods

### 1.2.1 Samples

Control and patient samples were collected after obtaining Institutional Ethics Committee approval for carrying out a study on biomarker discovery in plasma cell disorders. The work was carried out in accordance with the *Declaration of Helsinki* after obtaining written informed consent. Excess control serum samples were obtained from healthy blood product donors who provided a sample for preliminary screening. Patients with confirmed plasma cell disorders with monoclonal gammopathy detected by SPEP, serum IFE or FLC, or a combination of these tests were included for MALDI-TOF MS analysis using the extra serum sample.

### 1.2.2 SPEP, serum IFE and FLC Analysis

SPEP was performed on the Minicap/ FP fully automated capillary electrophoresis analyser (Sebia, France) and serum total protein concentration was determined by colorimetric method using biuret reagent on BA400 fully automated clinical chemistry analyser (manufactured by Bio system S.A., Spain) in the Department of Biochemistry. From the total protein value, the software automatically quantified each fraction (*α*1, *α*2, *β*1, *β*2 and γ globulins). With the help of a pointer on the screen, the M-band area was marked manually, and the instrument automatically calculated the concentration of M-protein from the total protein value. Serum IFE was performed on SAS-3 9IF gel (Helena Biosciences) and reported. sFLC was analysed on BN II Nephelometer by Siemens using the Binding Site kit.

### 1.2.3 Reagent-based extraction

The serum fraction was separated from whole blood by centrifugation at 5000 rpm for 15 minutes and stored at −80ΰC until further analysis. 25 *μL* of the serum was taken in a fresh 500 *μL* centrifuge tube and mixed with 400 *μL* of extraction solution (50% Acetonitrile (ACN) prepared in mass spectrometry grade water), the mixture was thoroughly mixed allowing the protein fraction to precipitate. Mass spectrometry grade ACN and LC-MS grade water were both obtained from Sigma-Aldrich. The mixture was incubated at room temperature (28ΰC) for 5 minutes. The mixture was centrifuged at 12,000 rpm for 10 minutes, and the protein precipitate was collected. The protein precipitate was washed with wash solution (20% ACN prepared in mass spectrometry grade water). It was mixed with equal volumes, i.e., 400 *μL* of 20% ACN and the precipitate was mixed thoroughly. The solution was centrifuged at 10,000 rpm for 10 minutes. The supernatant was discarded, and the precipitate was retained. After the wash, the wash solution was removed to the maximum extent possible. The precipitate was reconstituted in 200 *μL* reconstitution buffer: 10% formic acid (FA) and 50 mmol/L tris(2-carboxyethyl)phosphine hydrochloride (TCEP). The reconstituted solution was centrifuged at 12000 rpm for 10 minutes. TCEP was purchased from Tokyo Chemical Industry Co., LTD and formic acid with 98% purity purchased from Sigma-Aldrich.

### 1.2.4 Antibody Immuno-enrichment

The procedure for immunoenrichment was performed as previously described (7) with some modifications as described. Due to the non-availability of agarose beads and nanobodies targeting *κ* and *λ* constant region domains, our protocol was modified to include direct use of biotin-labelled *κ* and *λ* antibodies and subsequently immobilized in streptavidin magnetic beads. 25 *μL* of serum was diluted in 225 *μL* of LC-MS grade water and added to a mixture of 5 *μL* each of *κ* and *λ* antibodies along with 5 *μL* of streptavidin beads. Captureselect Biot A-light chain *κ* HU and light chain *λ* HU antibodies and Myone Streptavidin T1 beads were purchased from Thermo Fisher Scientific. The diluted serum, along with the antibody-coated bead mixture, were combined and incubated at room temperature for 45 minutes. This mixture was then washed five times with LC-MS grade water and eluted with 40 *μL* elution buffer (10% FA, 50 mmol/L TCEP). This was vortexed for 15 minutes.

### 1.2.5 MALDI-TOF Analysis

One *μL* of the protein solution was spotted along with MALDI matrix (alpha-cyano-4-hydroxycinnamic acid, 10 g/L in 50% Acetonitrile + 0.1% TFA) onto MALDI ground steel plate and allowed to dry. The samples were then analysed using Ultraflex™ LT, Bruker MALDI/TOF-TOF mass spectrometer. Mass measurement was analysed with a summation of 500-5000 shots depending on the intensity of the M-peak. The mass spectra for each sample was exported to FlexAnalysis 3.3 (Bruker Daltonics) and background subtracted. Before analysis, the MALDI mass spectrometer was calibrated using protein standard (Bruker Daltonics).

### 1.2.6 Interpretation of M-protein

A sample was considered positive for M-protein if there was a sharp or broad peak within the *κ* or *λ m/z* mass spectra range. (7) All the patient samples were overlaid on a healthy serum control mass spectra with the polyclonal distribution of *κ* and *λ* light chains.

## 1.3 Results

### 1.3.1 Identification of *κ* and *λ* chains in ACN precipitates of serum (Fig. 1a and b)

The amount of serum sample was tested in the range from 10 *μL*- 100 *μL*. 25 *μL* was chosen for further analysis because, at this loading amount, the intensity of M-peak made the detection unambiguous (data not shown). We tested a range of concentrations for ACN from 10% to 100% and found that 50% was optimal for analysis (data not shown). We spotted a range of sample concentrations on the MALDI plate: neat, 1:10, 1:25, 1:50 and 1:100. Neat and 1:10 concentration were found to be suitable for further analysis. The neat sample was spotted on the MALDI plate for all control and patient samples.

**Figure 1a:**
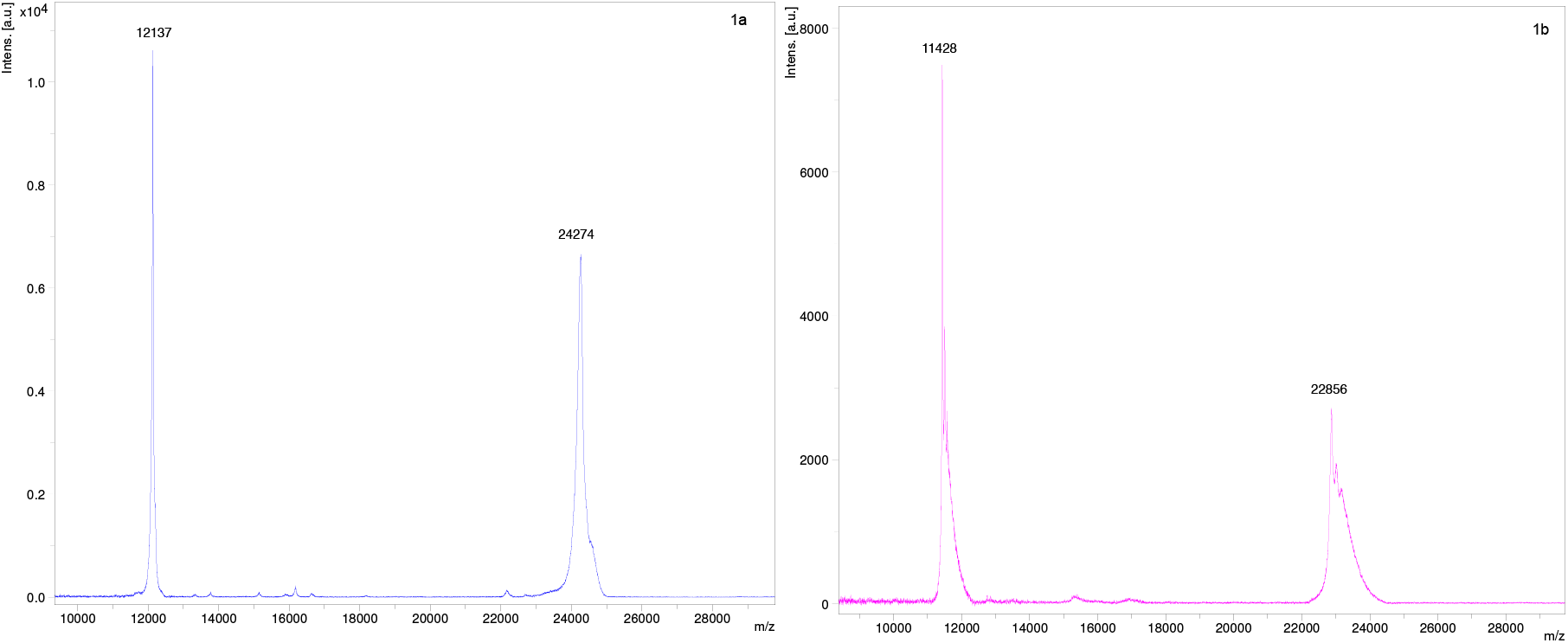
FLC mass spectra with [M+H]^+^ and [M+2H]^+2^ charge states for a patient serum (sample no.2-Table 1) with *κ* monoclonal gammopathy by Acetonitrile precipitation. [M+H]^+^−24274 Da & [M+2H]^+^−12137 Da. Figure 1b: FLC mass spectra with [M+H]^+^ and [M+2H]^+2^ charge states for a patient serum (sample no: 6) with *λ* monoclonal gammopathy by Acetonitrile precipitation. [M+H]^+^−22856 Da & [M+2H]^+^- 11428 Da.

All the images were acquired at a *m/z* range of 10000-29000 Da to include the [M+H]^+1^ and [M+2H]^+2^ charge states for *κ* and *λ*. [M+2H]^2+^charge state images were chosen for further analysis due to superior visual sensitivity as already described in the MASS-SCREEN analysis. (7)

**Table 1:** Baseline characteristics of patients and comparison of monoclonal gammopathy results by SPEP, serum IFE, sFLC and Acetonitrile precipitation by MALDI-TOF MS

| Sample No | Age and Sex | Diagnosis | ISS<sup>1</sup> Stage (for MM) | LDH<sup>2</sup> in (U/L) | SPEP: M-protein conc. (g/dL) | Serum IFE<sup>3</sup> | Serum free light chain<sup>4</sup> |  |  | M-peak (+2 charge state in Da) | MAL-TOF MS Analysis |
| --- | --- | --- | --- | --- | --- | --- | --- | --- | --- | --- | --- |
|  |  |  |  |  |  |  | Kappa (mg/L) | Lambda (mg/L) | Ratio |  |  |
| 1. | 67/F | MM | III | 393 | 4 | IgGκ | 75.5 | 27.5 | 2.745 | 11689 | κ |
| 2. | 50/F | MM | II | 421 | 3.3 | IgGκ | 655 | 5.87 | 111.584 | 12137 | κ |
| 3. | 63/F | Plasmacytoma | N/A<sup>5</sup> | N/A | 0.8 | IgGκ | N/D | N/D | N/D | 11725 | κ |
| 4. | 44/M | MM | I | 286 | 2.3 | IgGλ | 1.25 | 245 | 0.005 | 11489 | λ |
| 5. | 52/M | MM | II | Missed | 3.2 | IgGκ | 16.7 | 2.95 | 5.661 | 11869 | κ |
| 6. | 69/F | MM | III | 474 | 8.8 | IgGλ | 14.5 | 798 | 0.018 | 11428 | λ |
| 7. | 57/F | MM | II | 466 | 3.8 | IgGκ | 374 | 9 | 41.556 | 11681 | κ |
| 8. | 63/F | MM | III | 206 | 4 | IgAκ | 32.5 | 2.55 | 12.75 | 11678 | κ |
| 9. | 67/M | MM | II | 336 | 3 | IgGκ | 571 | 9.54 | 59.853 | 11708 | κ |
| 10. | 68/F | MM | III | 319 | Absent | λ | 34.6 | 2550 | 0.014 | 11488 | λ |
| 11. | 52/M | MM | III | 1022 | 4.9 | N/D<sup>6</sup> | N/D | N/D | N/D | 11720 | κ |
| 12. | 59/M | MM | II | 280 | 5 | N/D | 37.9 | 2.01 | 18.856 | 11636 | κ |
| 13. | 54/F | Plasmacytoma | N/A | N/A | 1.2 | N/D | N/D | N/D | N/D | 11751 | κ |
| 14. | 58/M | MM | III | 260 | 4.7 | IgGκ | N/D | N/D | N/D | 11692 | κ |
| 15. | 47/M | MM | II | 260 | 2.8 | IgAκ | 90.1 | <0.05 | Not calculable | 11558 | κ |
| 16. | 48/F | MM | II | 445 | Absent | κ | 6070 | 3.3 | 1839 | 11797 | κ |
| 17. | 64/M | MM | N/A | N/A | Absent | Normal | 1433 | 13.79 | 103 | 11796 | κ |
| 18. | 59/F | Plasmacytoma | N/A | N/A | Absent | N/D | 51 | 16 | 3.188 | 11818 | κ |
| 19. | 46/F | MM | II | 830 | 2.2 | IgGλ | 1.17 | 1530 | 0.001 | 11377 | κ |
| 20. | 44/M | MM | I | 415 | 1.2 | IgGλ | 16.9 | 16.2 | 1.043 | 11649 | κ |
| 21. | 62/F | MM | III | 596 | 1.9 | IgAκ | 3710 | 9.06 | 409.492 | 11835 | κ |
| 22. | 72/F | MM | II | 283 | 2.2 | IgGκ | 2740 | 9.31 | 294.307 | 11692 | κ |
| 23. | 63/M | MM | III | 1657 | 2.7 | IgAκ | 1460 | 10.2 | 143.14 | 12128 | κ |
| 24. | 70/M | MM | I | 330 | Absent | κ | 3980 | 12.2 | 326.23 | 11729 | κ |
| 25. | 64/M | MM | II | 379 | 2.3 | N/D | 61.3 | 18.7 | 3.278 | 11706 | κ |
| 26. | 64/F | MM | II | 200 | 4.2 | IgGκ | 70.3 | 3.27 | 21.5 | 11772 | κ |
| 27. | 63/F | MM | II | 340 | 2.8 | IgGκ | 3750 | 8.39 | 446 | 11580 | κ |

<sup>1</sup> ISS- International staging system for multiple myeloma: I-serum β2 microglobulin <3.5 mg/L and serum albumin ≥3.5 g/dL; II- Not stage
I or III; III- serum β2 microglobulin ≥5.5 mg/L
<sup>2</sup> LDH normal range: 200-400 U/L (serum LDH is a prognostic marker in the R-ISS staging system)
<sup>3</sup> IFE was reported at Dr LalPath labs (IgD and IgE immunotyping was not performed)
<sup>4</sup> Serum free light chain was reported at SRL Diagnostics
<sup>5</sup> N/A- Not applicable
<sup>6</sup> N/D- Not done (missing)

### 1.3.2 Validation of ACN precipitation results with Immunoenrichment with *κ* (Fig. 2a and b) and *λ* antibodies (Fig. 3a and b)

Immunoenrichment was performed for two known *κ* M-protein samples and, two *λ* M-protein samples. The mass spectra were found to be identical with the light chain *m/z* falling within the range as reported by immunoenrichment.

**Figure 2:**
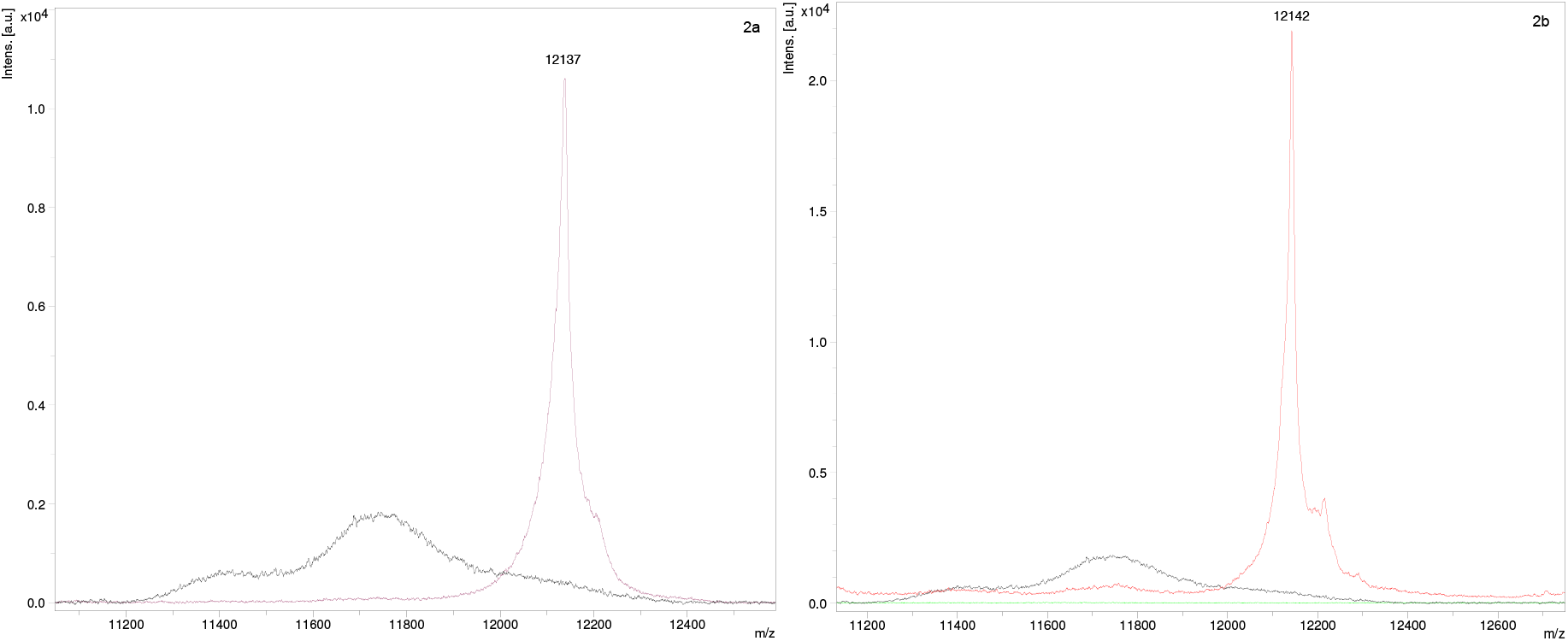
Patients serum (sample no. 2-Table 1) sample with *κ* monoclonal gammopathy Figure 2a: FLC mass spectra showing a sharp M-peak (violet) overlaid on a healthy serum control (black) by Acetonitrile precipitation. Figure 2b: Immunoenrichment with *κ* antibody tagged to streptavidin beads from the same patient showing a sharp M-peak (red) overlaid over a healthy serum control (black) and streptavidin bead control with no peak (green).

**Figure 3:**
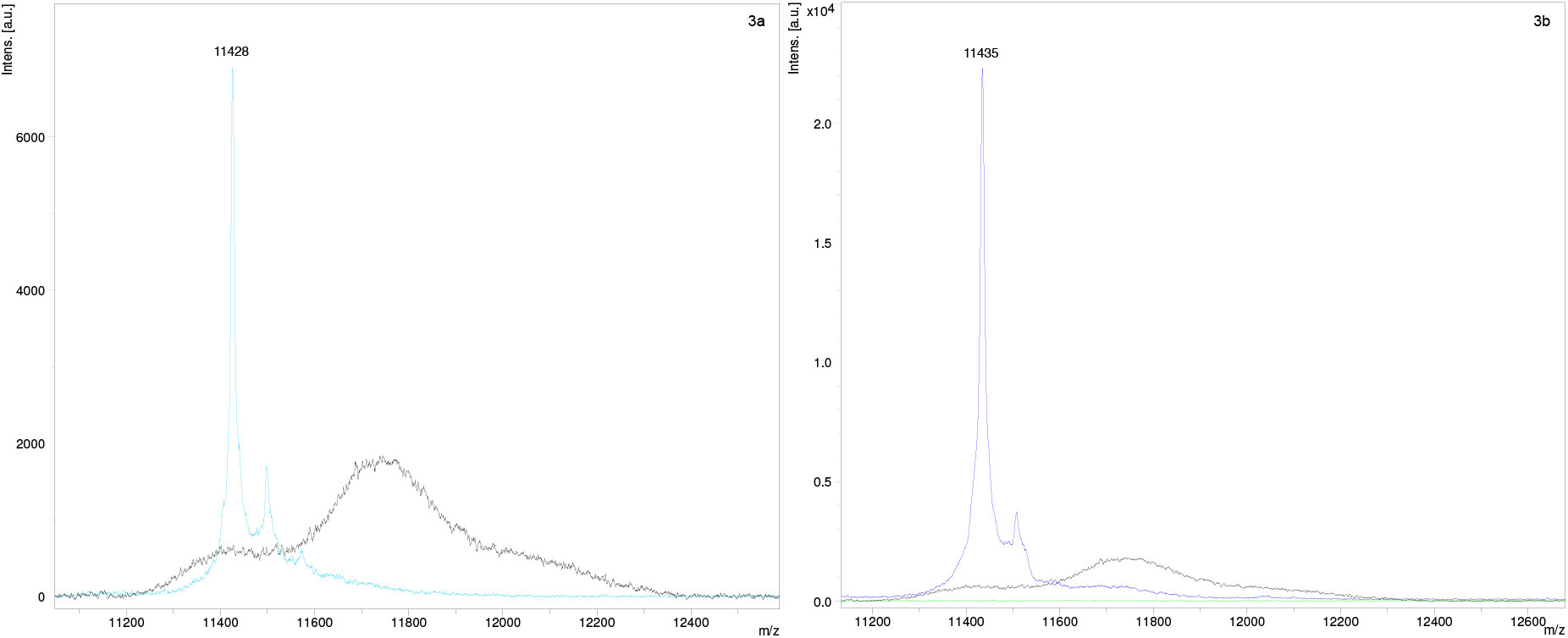
Patient’s serum (sample no. 6-Table 1) with *λ* monoclonal gammopathy Figure 3a: FLC mass spectra showing a sharp M-peak (blue) overlaid on a healthy serum control (black) by Acetonitrile precipitation. Figure 3b: Immunoenrichment with *λ* antibody tagged to streptavidin beads from the same patient showing a sharp M-peak (blue) overlaid over a healthy serum control (black) and streptavidin bead control with no peak (green).

### 1.3.3 ACN precipitation of the control serum samples (Fig. 4)

The Gaussian distribution of light chains was obtained by analysing 20 control samples from extra serum samples of apparently healthy blood donors. All the control samples underwent SPEP. Serum protein concentration for all samples was within normal limits, and SPEP did not demonstrate M-protein.

**Figure 4:**
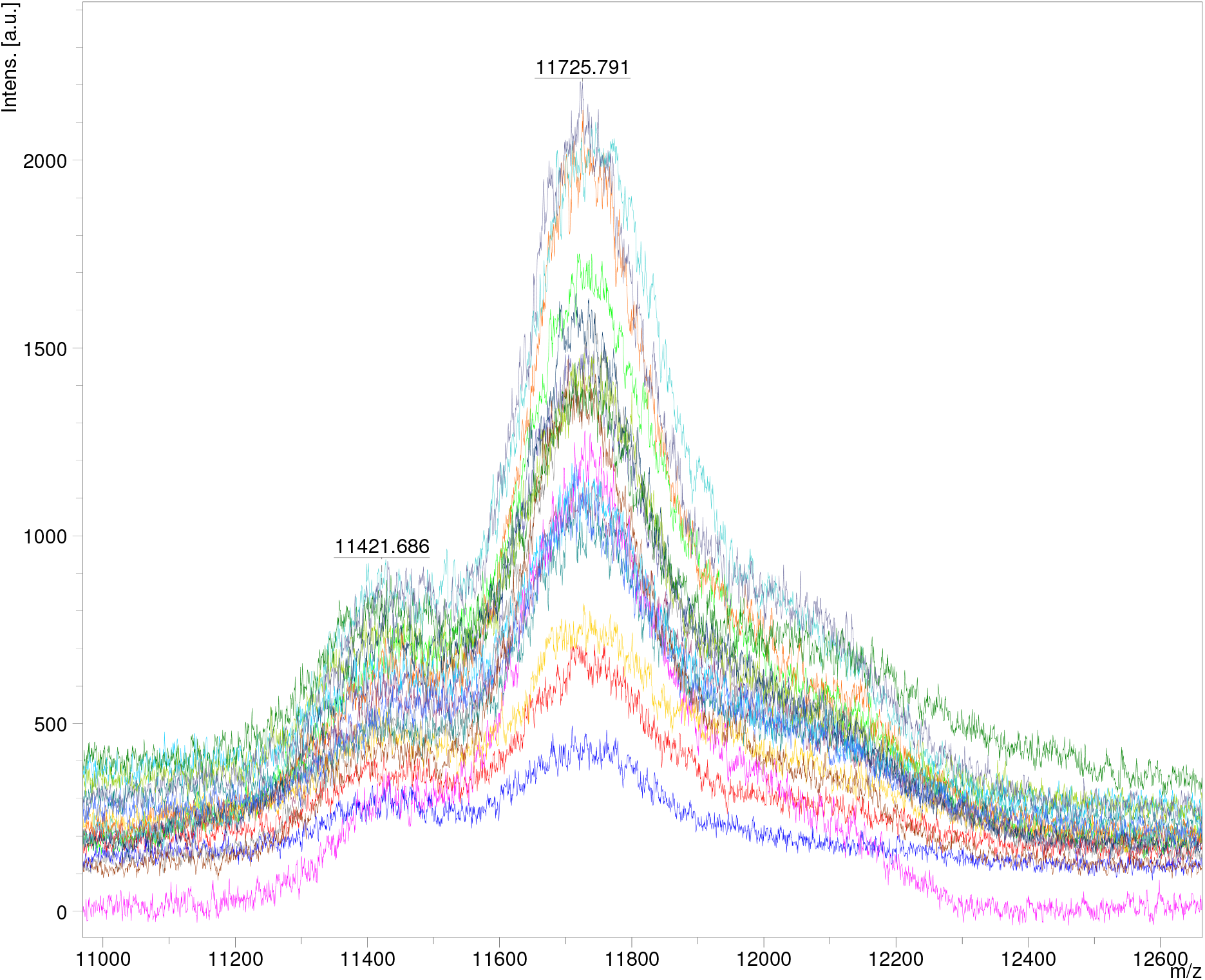
Mass spectra at [M+2H]^2+^ charge state of all the healthy serum controls demonstrating the Gaussian distribution of polyclonal *κ* & *λ* light chains.

### 1.3.4 Patient samples

Twenty-four patients were newly diagnosed with multiple myeloma and, three with solitary plasmacytoma. The baseline characteristics of these patients are shown in Table 1.

A mass spectrometrist S.J was blinded to the IFE and FLC results-blinded analyst. N.M was the unblinded analyst. All the 27 samples with monoclonal gammopathy confirmed by the other biochemical techniques, demonstrated a positive M-peak on MALDI-TOF MS (100%).

### 1.3.5 Confounding samples (Fig. 5)

Three samples were labelled as confounders due to low peak intensity in the *κ* mass spectra. However, their peaks matched the sFLC results as reported on serum IFE and sFLC.

**Figure 5:**
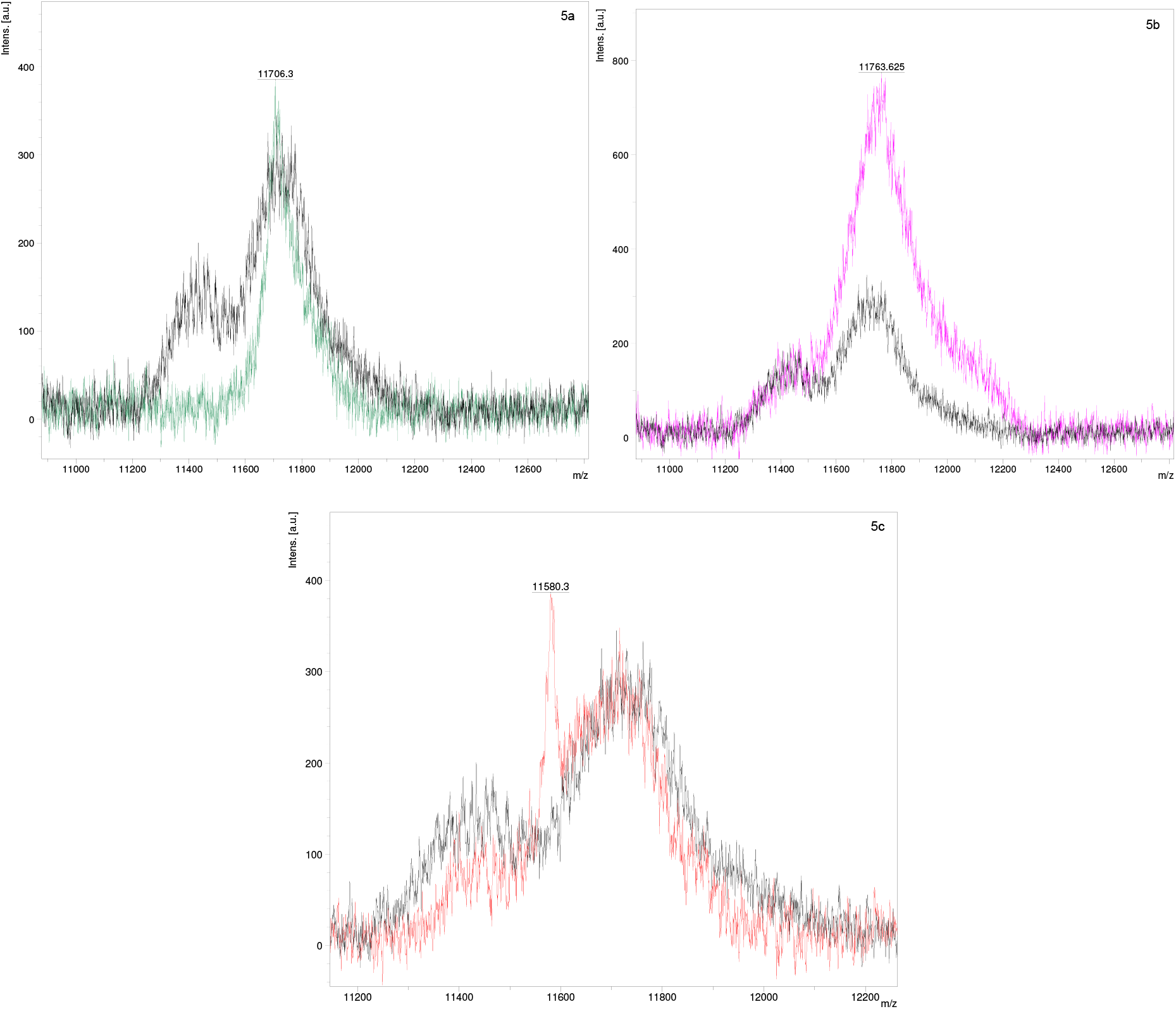
Mass spectra of the confounding samples-5a and b: Serum (sample no. 25 and 26, respectively-Table 1) from two different patients with *κ* monoclonal gammopathy showing low intensity (green and violet, respectively) in comparison with a healthy serum control (black). Note: The *m/z* of the peak in these patient’s samples falls within the *κ m/z* range. 5c: Mass spectra of serum (sample no. 27-Table 1) from a patient with *κ* monoclonal gammopathy confirmed by serum IFE and sFLC analysis showing a low-intensity *κ* peak (red) overlaid on a healthy serum control (black).

### 1.3.6 Concordance with IFE

Serum IFE and sFLC analysis were sent to independent laboratories for external validation of the MALDI-TOF MS results.

Among all the 27 patients with positive M-protein on MALDI-TOF MS, 23 patients had serum IFE results available. Of the 23 patients, IFE and MALDI-TOF MS reports were concordant in 21 patients (91%).

One patient (sample no: 20-Table 1) had IgG*λ* reported by serum IFE and sFLC was within normal limits. MALDI-TOF MS demonstrated a clear *κ* M-protein. Immunoenrichment was done which confirmed *κ m/z* M-protein. Another patient (sample no: 17-Table 1), had very high *κ* levels reported on sFLC: *κ* −1433 mg/L; *λ* −13.79; *κ*: *λ* ratio-103. However, serum IFE was reported normal (repeated twice in two different labs). This patient’s MALDI-TOF MS revealed a *κ* M-peak. Therefore, the accuracy in determining the FLC by IFE versus MALDI-TOF MS was 91%. Accuracy can be given for patient sample no: 20 (Table 1), but here the sFLC was within normal limits, indicating that MALDI-TOF MS was correct in 21/21.

### 1.3.7 Concordance with sFLC

Among all the 27 patients with positive M-protein on MALDI-TOF MS, 24 patients underwent sFLC testing. Of the 24 patients, 23 patients (96%) had concordant results on MALDI-TOF and sFLC demonstrating an altered *κ*/ *λ* ratio (>1.65 for *κ* or <0.26 for *λ*). One patient had a normal sFLC report- *κ*-19.9 mg/L; *λ*-16.2 mg/L; ratio- 1.043. IFE, MALDI-TOF MS and immunoenrichment confirmed *κ* M-protein on this patient sample. Therefore, the false-negativity rate of FLC determination by nephelometry analysis of sFLC was 4%.

## 1.4 Discussion

This study demonstrates the feasibility of qualitatively identifying M-protein without the need for nanobody immunoenrichment using agarose or sepharose beads. (12).

M-peak was accurately identified in 100% of patient serum samples by ACN precipitation using MALDI-TOF MS; in three samples there were low-intensity *κ* peaks which correlated with the free light chain detected on serum IFE and sFLC.

This technique demonstrates proof of principle of identifying monoclonal and polyclonal *κ* and *λ* as an alternative methodology for the demonstration of *m/z* mass spectra by MALDI-TOF MS. The results of this can be further verified by immunoenrichment.

The results of this study establish the ability to identify the FLC pattern correctly by ACN precipitation using MALDI-TOF MS in all patient serum samples. MALDI-TOF MS performed better than the other biochemical assays in the qualitative determination of FLC M-protein.

The drawbacks of this study include the small sample size. However, our results confirm the findings published previously regarding the superior analytical sensitivity of M-protein analysis by MALDI-TOF MS. (6–8) There is a need to analyse further the low M-peak intensity identified in the confounding samples which have not been addressed in the present study. The ACN precipitation technique was a qualitative analysis of M-protein as defined by its light chain clone. Further studies will focus on developing a methodology for the analysis of immunoglobulin isoforms and quantitative determination of M-protein using this methodology, as already reported with nanobody enrichment-MASS-FIX. (6) This methodology can also be utilised for the estimation of M-protein in the urine.

The ACN precipitation methodology, a low-cost technique could be utilised for its cost-effectiveness in planning an epidemiological study in MGUS in a geographically-defined population in a low and middle-income country like India. This technique can also serve as a useful screening technique for M-protein in a haematology lab. MALDI-TOF MS has also shown an advantage of its use in minimal residual disease (MRD) assessment when compared to bone marrow-based MRD analysis by flow cytometry. (13)

## 1.5 Conclusions

We have developed a low-cost reagent-based extraction process using Acetonitrile precipitation to enrich for *κ* and *λ*, for the screening and qualitative analysis of M-protein.

## Abbreviations

MALDI-TOF: matrix-assisted laser desorption ionization-time of flight
MS: mass spectrometry
SPEP: serum protein electrophoresis
IFE: Immunofixation electrophoresis
sFLC: Serum free light chains
ACN: Acetonitrile
MM: multiple myeloma
*κ*: Kappa
*λ*: Lambda
*m/z*: Mass to charge ratio

## 1.6 Acknowledgements

We acknowledge the Institute’s blood bank staff for helping in the collection of control serum samples for the study, the residents of the Department of Medical Oncology and the Biochemistry staff for coordinating the collection of patient samples.

