## Supplemental material for "Detection of M-Protein in Acetonitrile Precipitates of Serum using MALDI-TOF Mass Spectrometry"

### Supplementary material

Figure 1- Mass spectra of patient sample no.1 with κ monoclonal gammopathy by MALDI-TOF MS overlaid on a healthy control

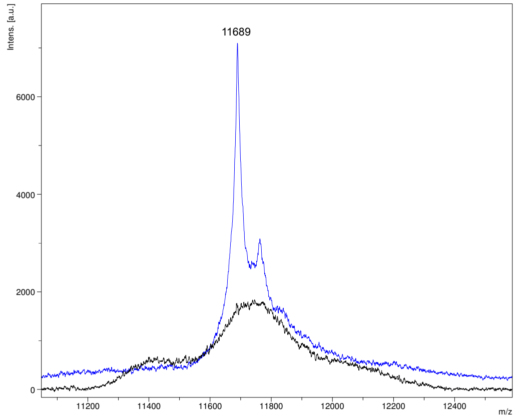

Figure 2- Mass spectra of patient sample no: 3 with κ monoclonal gammopathy by MALDI-TOF MS overlaid on a healthy control

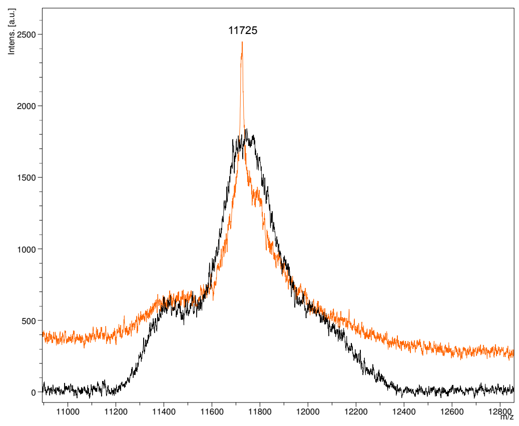

Figure 3- Mass spectra of patient sample no: 4 with λ monoclonal gammopathy by MALDI-TOF MS overlaid on a healthy control

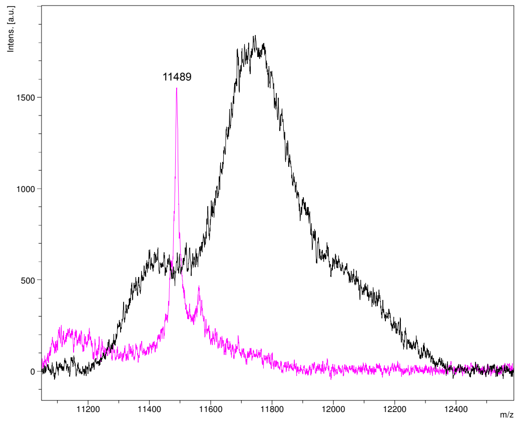

Figure 4- Mass spectra of patient sample no: 5 with κ monoclonal gammopathy by MALDI-TOF MS overlaid on a healthy control

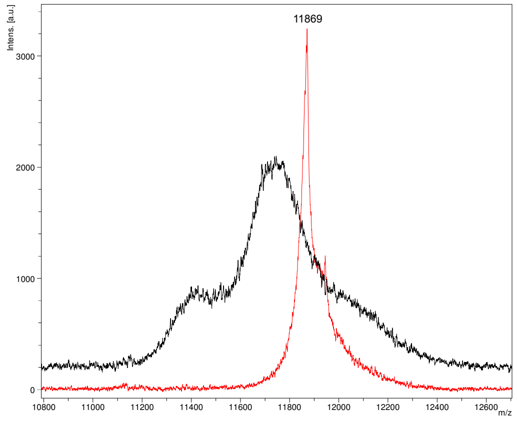

Figure 5- Mass spectra of patient sample no: 7 with $\kappa$ monoclonal gammopathy by MALDI-TOF MS overlaid on a healthy control

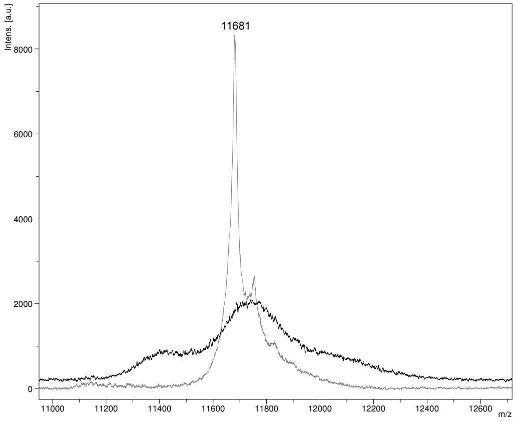

Figure 6- Mass spectra of patient sample no: 8 with κ monoclonal gammopathy by MALDI-TOF MS overlaid on a healthy control

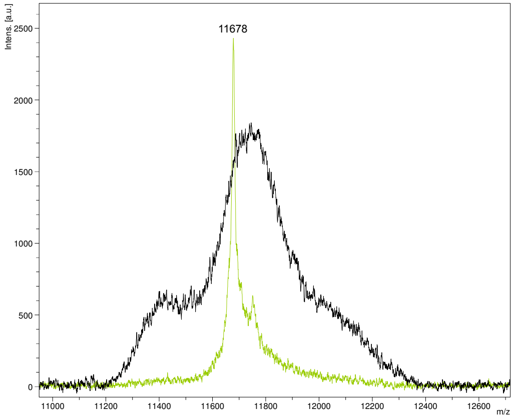

Figure 7- Mass spectra of patient sample no: 9 with $\kappa$ monoclonal gammopathy by MALDI-TOF MS overlaid on a healthy control

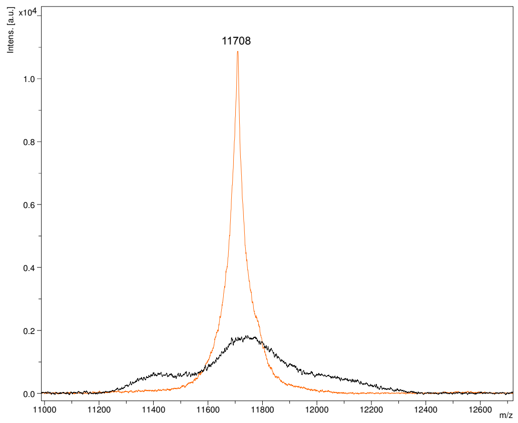

Figure 8- Mass spectra of patient sample no: 10 with λ monoclonal gammopathy by MALDI-TOF MS overlaid on a healthy control

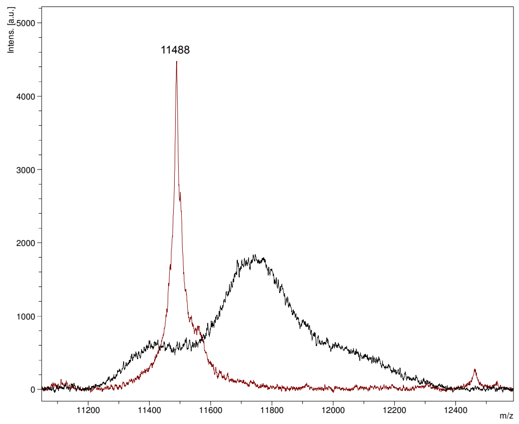

Figure 9- Mass spectra of patient sample no: 11 with κ monoclonal gammopathy by MALDI-TOF MS overlaid on a healthy control

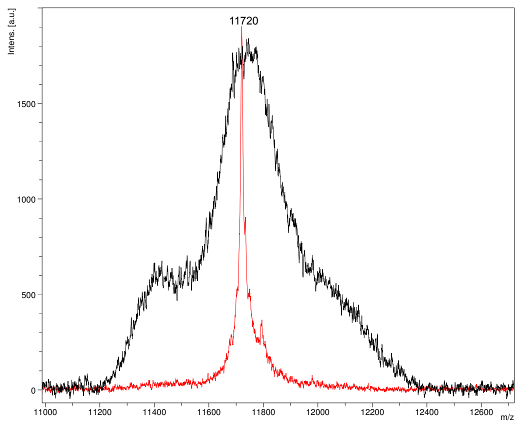

Figure 10- Mass spectra of patient sample no: 12 with κ monoclonal gammopathy by MALDI-TOF MS overlaid on a healthy control

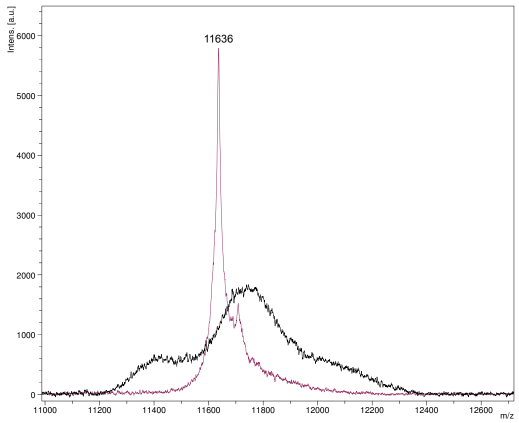

Figure 11- Mass spectra of patient sample no: 13 with κ monoclonal gammopathy by MALDI-TOF MS overlaid on a healthy control

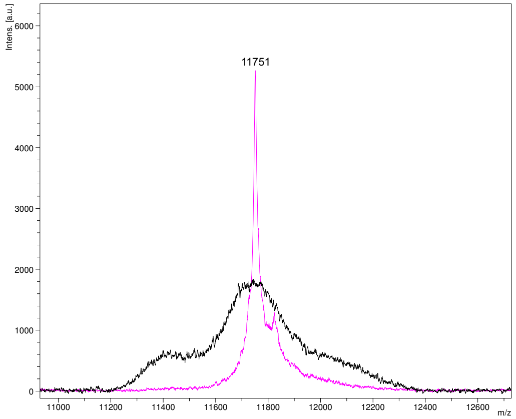

Figure 12- Mass spectra of patient sample no: 14 with κ monoclonal gammopathy by MALDI-TOF MS overlaid on a healthy control

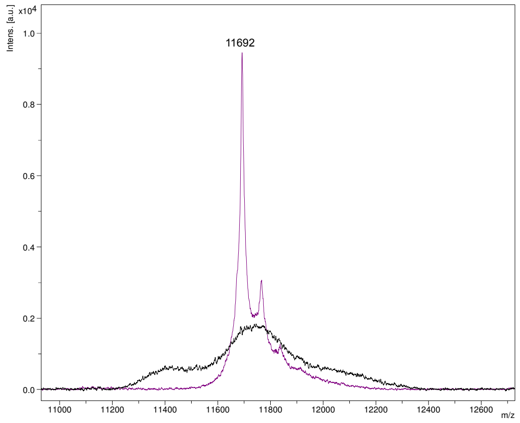

Figure 13- Mass spectra of patient sample no: 15 with κ monoclonal gammopathy by MALDI-TOF MS overlaid on a healthy control

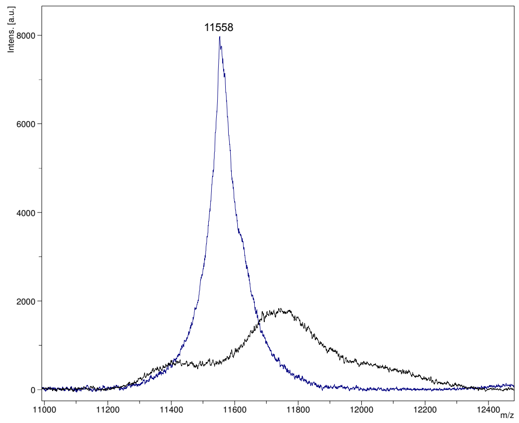

Figure 14- Mass spectra of patient sample no: 16 with $\kappa$ monoclonal gammopathy by MALDI-TOF MS overlaid on a healthy control

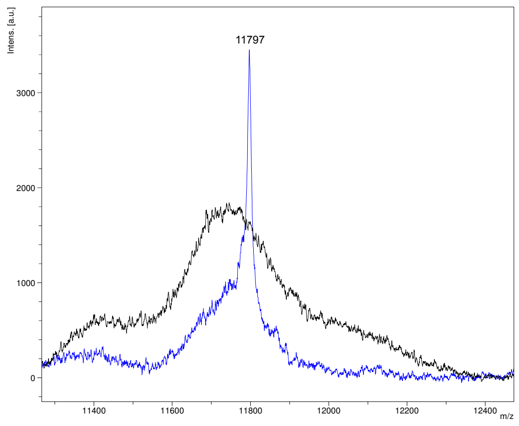

Figure 15- Mass spectra of patient sample no: 17 with $\kappa$ monoclonal gammopathy by MALDI-TOF MS overlaid on a healthy control

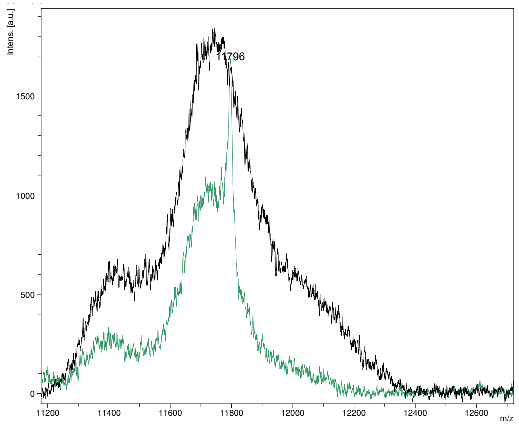

Figure 16- Mass spectra of patient sample no: 18 with κ monoclonal gammopathy by MALDI-TOF MS overlaid on a healthy control

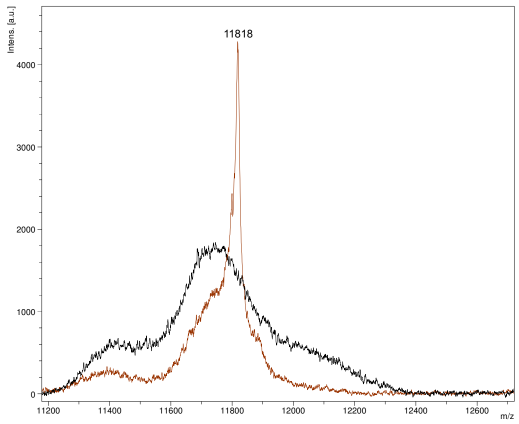

Figure 17- Mass spectra of patient sample no: 19 with κ monoclonal gammopathy by MALDI-TOF MS overlaid on a healthy control

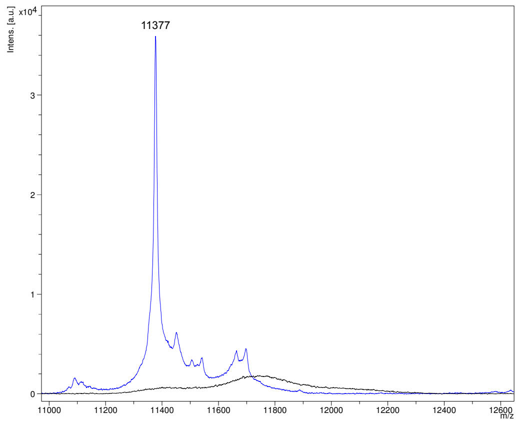

Figure 18- Mass spectra of patient sample no: 20 with $\kappa$ monoclonal gammopathy by MALDI-TOF MS overlaid on a healthy control

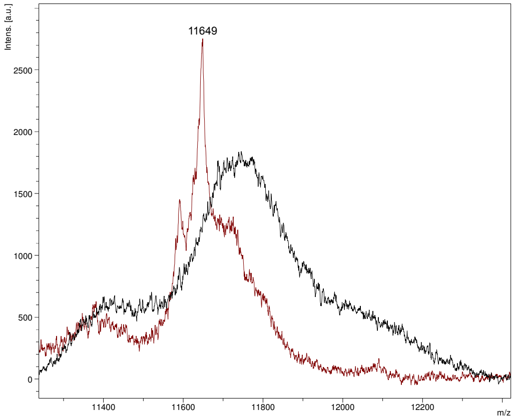

Figure 19- Mass spectra of patient sample no: 21 with κ monoclonal gammopathy by MALDI-TOF MS overlaid on a healthy control

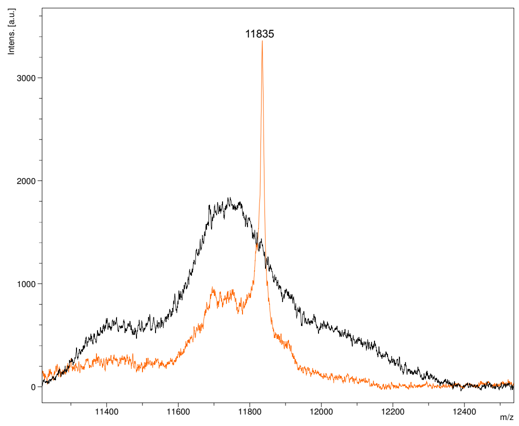

Figure 20- Mass spectra of patient sample no: 22 with $\kappa$ monoclonal gammopathy by MALDI-TOF MS overlaid on a healthy control

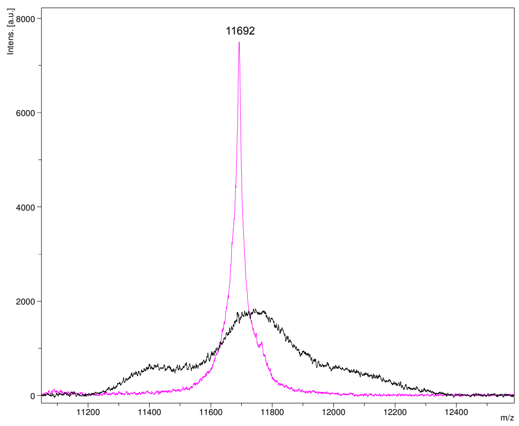

Figure 21- Mass spectra of patient sample no: 23 with $\kappa$ monoclonal gammopathy by MALDI-TOF MS overlaid on a healthy control

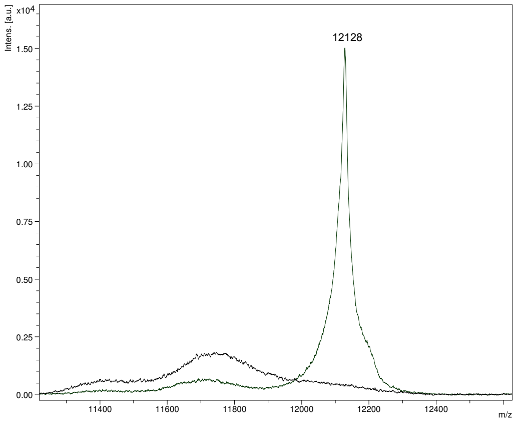

Figure 22- Mass spectra of patient sample no: 24 with κ monoclonal gammopathy by MALDI-TOF MS overlaid on a healthy control

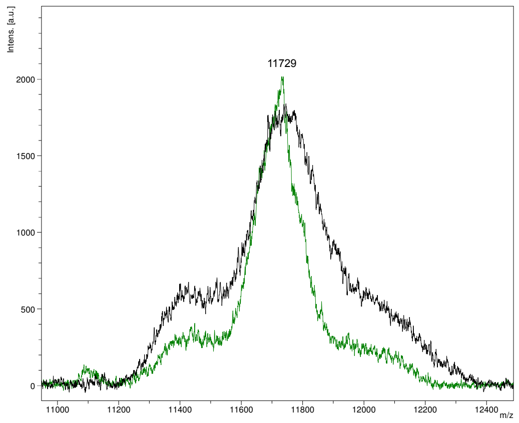
